# Annual and 16-day rangeland production estimates for the western United States

**DOI:** 10.1101/2020.11.06.343038

**Authors:** Matthew O. Jones, Nathaniel P. Robinson, David E. Naugle, Jeremy D. Maestas, Matthew C. Reeves, Robert W. Lankston, Brady W. Allred

## Abstract

Rangeland production is a foundational ecosystem service and resource upon which livestock, wildlife, and people depend. Capitalizing on recent advancements in the use of remote sensing data across rangelands we provide estimates of herbaceous rangeland production from 1986-2019 at 16-day and annual time steps and 30m resolution across the western United States. A factorial comparison of this dataset and three national scale datasets is presented, and we highlight a multiple lines of evidence approach when using production estimates in decision-making. Herbaceous aboveground biomass at this scale and resolution provides critical information applicable for management and decision-making, particularly in the face of annual grass invasion and woody encroachment of rangeland systems. These readily available data remove analytical and technological barriers allowing immediate utilization for monitoring and management.

## Introduction

Rangeland production–specifically herbaceous aboveground biomass–is a foundational ecosystem service upon which livestock, wildlife, and people depend. Estimates of production have long been available via field-based measurements, but such estimates are geographically and temporally limited. Although statistical sampling techniques employed by national monitoring programs (Herrick et al., 2017; MacKinnon et al., 2011) provide means to monitor production across rangelands, such techniques and programs do not capture temporal variability or spatial heterogeneity at scales relevant to management and decision-making (e.g. within or across management units), with limited field-based plots often extrapolated to ecoregion scales (Karl et al., 2016). Satellite and airborne remote sensing methods (Smith et al., 2019) informed by field-based data can provide spatially contiguous and temporally continuous estimates of rangeland production. Methodologies often use empirical relationships between remote sensing indices (e.g. Normalized Difference Vegetation Index; NDVI) or terrestrial lidar retrievals, and measured or estimated biomass from field plots to map production. This methodology is most often applied at local or regional spatial scales (Jansen et al., 2018), but has also been successfully implemented at broad national scales (Reeves et al., 2020).

Vegetation production may also be estimated using remotely sensed data in process-based models, such as a light use efficiency model (Monteith, 1972) that calculates gross or net primary production (GPP and NPP, respectively) based on remotely sensed estimates of absorbed photosynthetically active radiation, the biophysical properties of vegetation types, and water and temperature constraints. These GPP and NPP models are prolific (Clark et al., 2011; Running et al., 2004) but the common units of carbon (i.e., g C m^−2^ year^−1^) are not relevant to rangeland managers or practitioners. These models also require land cover data which until recently were categorical at the pixel scale for U.S. rangelands and produced at 5-year time steps (Homer et al., 2015). Models could therefore not account for within-pixel heterogeneity of rangeland plant functional types (PFT) (e.g., annual grasses and forbs, perennial grasses and forbs, shrubs, or trees) or PFT variation in phenology and productivity (Browning et al., 2019). Accounting for this heterogeneity is especially critical considering the prominence of large-scale woody encroachment and annual grass invasion into rangeland systems (Jones et al., 2020). Continuous land cover datasets at annual time steps are now available (Allred et al., 2021, Rigge et al., 2020) allowing models to quantify changes in production in response to encroachment and invasion at temporal intervals relevant to management.

This technical note details the development of annual and 16-day rangeland herbaceous production estimates partitioned to perennial grasses/forbs and annual grasses/forbs across western U.S. rangelands. The data are easily accessible, account for within-pixel vegetation PFT heterogeneity, and are provided at temporal and spatial resolutions (and units) relevant to management. We perform a factorial comparison of this new production dataset and three national scale datasets. We highlight the value of using all data in a ‘multiple lines of evidence’ approach when implementing production estimates, where incorporating data derived from different methods into a decision-making process can spur greater data acceptance and application, and advance conservation of this valuable rangeland resource.

## Methods

### Net primary production partitioning

Detailed descriptions of the method used to calculate NPP by PFT are provided in Robinson et al., (2019); we provide an overview here. We produced spatially contiguous 16-day Landsat NDVI composites (Robinson et al., 2017) using Landsat 5 TM, 7 ETM+, and 8 OLI surface reflectance (Vermote et al., 2016) from 1986-2019 across Western U.S. rangelands (Reeves and Mitchell, 2011). Using the 16-day NDVI and a PFT cover dataset (Allred et al., 2021), we disaggregated pixel level NDVI using linear mixing theory to its sub-pixel PFT components. In brief, the NDVI of a mixed pixel is disaggregated to the PFTs present in the pixel, weighted by their fractional cover and the ecoregion-scale phenology of each PFT; the mean of the PFT specific NDVI values equate to the mixed pixel NDVI. To capture and incorporate the geographically specific PFT NDVI phenological characteristics, we built an overdetermined set of linear equations (Robinson et al., 2019) to solve for each PFT NDVI value within US EPA Level IV regions (Omernik and Griffith, 2014). The result is PFT NDVI estimations that capture NDVI amplitudes and regional PFT phenology. We reprojected and bilinearly resampled all Landsat imagery to a geographic coordinate system of approximately 30m resolution prior to manipulation.

The PFT specific NDVI values are then used in the MOD17 NPP model adapted for Landsat (Robinson et al., 2018). Using linear interpolation we calculated daily NDVI values between each 16-day composite. We then calculated daily NPP for each PFT present in the pixel using the daily PFT NDVI values, daily GRIDMET meteorology (Abatzoglou, 2013), and the specific PFT’s biophysical properties (Robinson et al., 2018). We multiplied NPP estimates by the PFT fractional cover estimates, resulting in total grams of carbon assimilated per PFT per pixel per day (g C m^−2^ day^−1^). Daily values are summed to 16-day values (g C m^−2^ 16days^−1^) and to annual values (g C m^−2^ year^−1^).

### NPP conversion to herbaceous aboveground biomass

The herbaceous NPP, partitioned to perennial grasses/forbs and annual grasses/forbs, is allocated to aboveground (ANPP) pools using the equations:

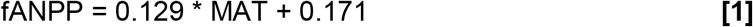

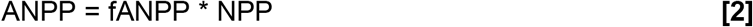

where fANPP is the fraction partitioned to ANPP and MAT is mean annual temperature (Hui and Jackson, 2006). We convert ANPP (g C m^−2^ 16 days^−1^) to biomass (kg ha^−1^ or lbs acre^−1^) using the pixel area and a 47.5% carbon content of vegetation estimate (Eggleston et al., 2006); the midpoint of a 45% to 50% carbon to dry matter estimation range (Schlesinger, 2005).

### Comparisons

We calculated Pearson correlation coefficients between the herbaceous aboveground biomass (HAGB) estimates and 16,591 Natural Resources Conservation Service (NRCS) National Resources Inventory (NRI) plot-level estimates of herbaceous biomass collected on rangelands from 2004 to 2018 (NRCS, USDA, 2015). The HAGB estimate corresponding to each plot was sampled from the same year as the plot measurement.

We also compared HAGB estimates to United States Forest Service Rangeland Production Monitoring Service (RPMS) data, provided annually from 1984-2018 at 250m resolution (Reeves et al., 2020), and to the gridded Soil Survey Geographic (gSSURGO) database which provides fixed estimates of unfavorable, normal, and favorable annual range potential production by soil survey units at 30m resolution (Soil Survey Staff, 2017). The RPMS and gSSURGO data estimate total rangeland productivity (not solely herbaceous) but are used here as the only available gridded productivity datasets that are specific to western U.S. rangeland systems and cover a similar time period. To account for temporal variability, we compared the 50th percentile of the two temporally dynamic datasets using years 2000-2018, and the gSSURGO ‘normal’ data. We calculated Pearson correlation coefficients using a subsample (5000 random rangeland locations) of each of the three datasets (RAP HAGB 50th percentile, RPMS 50th percentile, gSSURGO normal). For all comparisons we only included rangelands identified by Reeves and Mitchell (2011) inclusive of afforested, pasture, and barren categories. We also calculated the difference between the gridded HAGB, RPMS, and gSSURGO estimates and the plot level NRI herbaceous biomass estimates; scatterplots, correlations, and the geographic distribution of those differences are provided in supplemental information.

## Results

### Herbaceous aboveground biomass

Estimates of HAGB at 30m resolution are provided annually (Fig. 1a-c) from 1986-2019, and as accumulating HAGB at 16-day intervals (Fig. 1d). The HAGB is partitioned into perennial grasses/forbs and annual grasses/forbs and accounts for variation in pixel-scale fractional cover at annual time steps (Allred et al., 2021) and the phenology of each PFT. The data are accessible for viewing and analysis via a publicly available online application, the Rangeland Analysis Platform (https://rangelands.app/).

**Figure 1.**
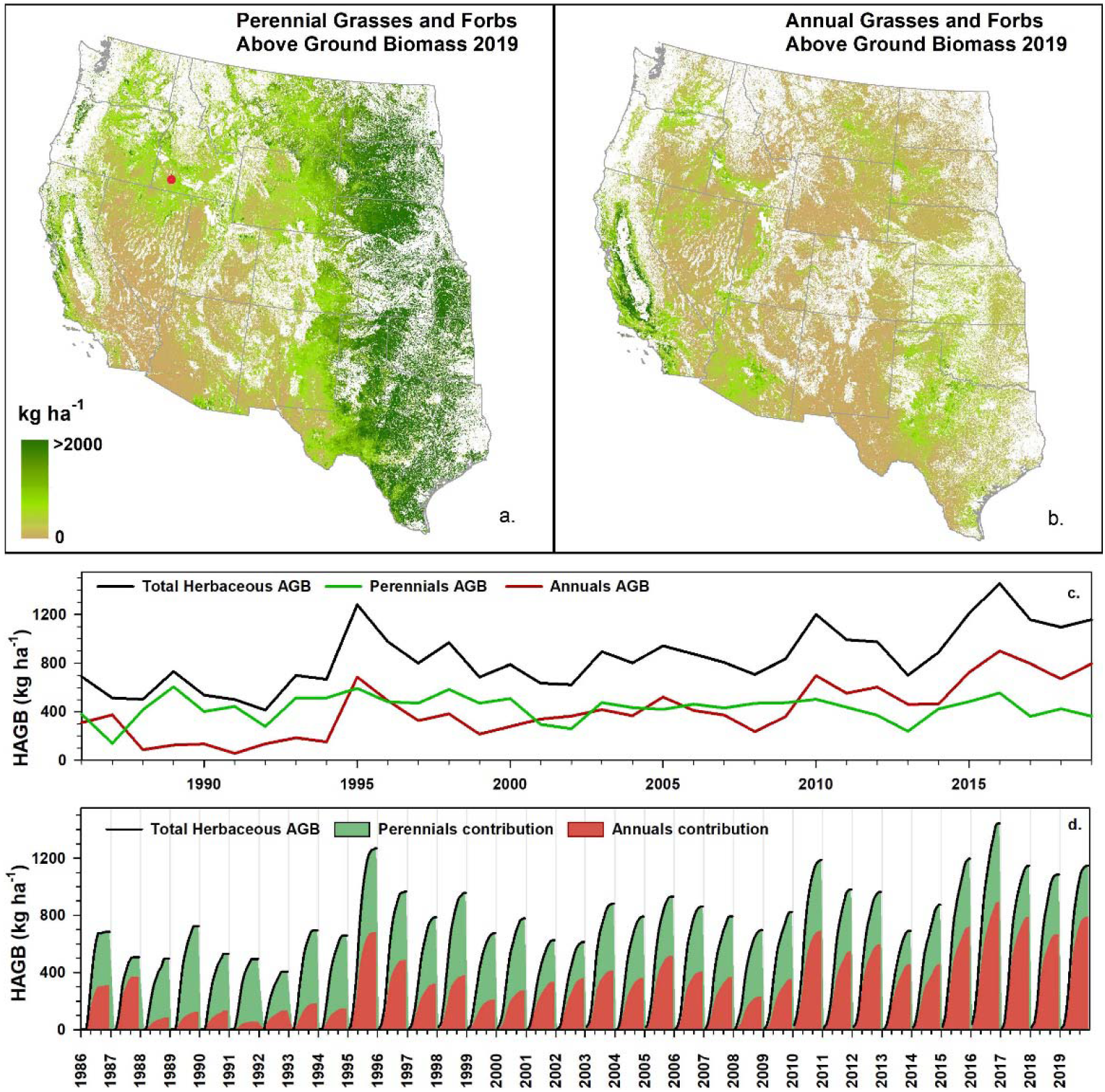
Herbaceous aboveground biomass (HAGB) at 30m resolution across western U.S. rangelands available via the Rangeland Analysis Platform (https://rangelands.app/). Total annual 2019 HAGB partitioned to (a) perennial grasses and forbs and (b) annual grasses and forbs. (c) Total yearly HAGB and portions attributable to perennials and annuals for a Bureau of Land Management grazing allotment in southwest Idaho (red point in a.) from 1986-2019. (d) For the same grazing allotment, 16-day accumulating total HAGB and partitioned contributions from perennial and annual grasses/forbs for 1986-2019.

### Data comparisons

The HAGB data available via the Rangeland Analysis Platform (hereafter RAP HAGB) are well correlated (r = 0.63) with 16,591 NRI plot-level herbaceous biomass estimates (Fig. 2).

**Figure 2.**
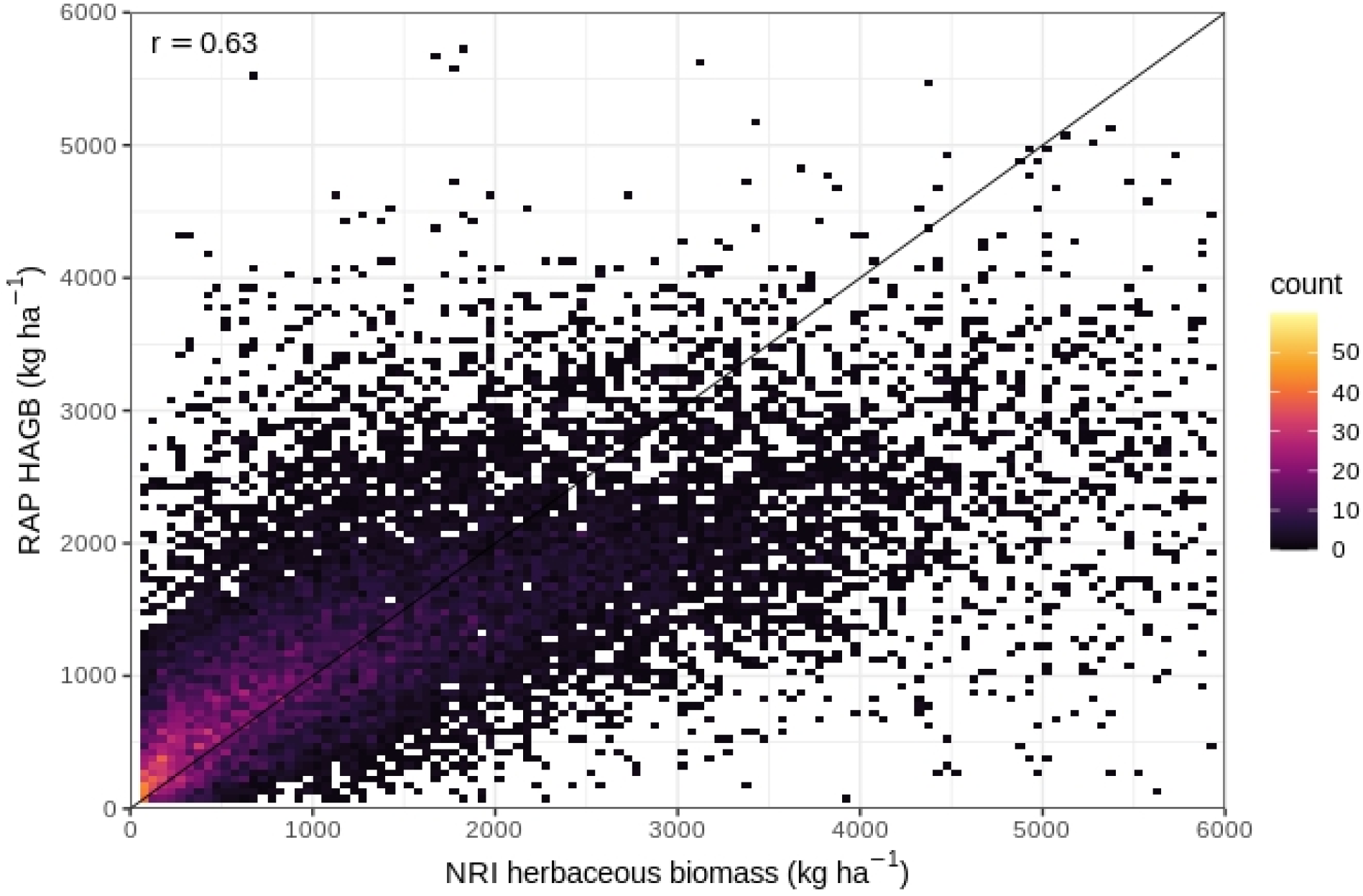
Density scatterplot (bin width 50 kg ha^−1^) of RAP herbaceous aboveground biomass (HAGB) and 16,591 NRI plot-level biomass estimates, 1:1 line (black), and Pearson correlation coefficient (r=0.63). NRI biomass estimates greater than 6000 kg ha^−1^ are not displayed.

We subtracted the RPMS 50th percentile and gSSURGO ‘normal’ gridded data from the RAP HAGB 50th percentile (as well as the gSSURGO from RPMS) which provided the geospatial distribution of differences between the products. Mapped differences (Fig. 3a-c) and scatterplots of 5000 randomly sampled rangeland locations (Fig. 3d-f) display strong agreement as indicated by Pearson correlation coefficients. At lower biomass levels (< ~1500 kg ha^−1^) the RAP HAGB 50th percentile is well aligned with the RPMS HAGB 50th percentile (Fig. 3d) and the gSSURGO normal (Fig. 3e). The RAP HAGB displays generally lower estimates than the other two data sets at higher biomass levels while RPMS and gSSURGO are more evenly distributed along the 1:1 line (Fig. 3f). These distributions are expected as RPMS and gSSURGO provide total production estimates while the RAP HAGB is herbaceous production only.

**Figure 3.**
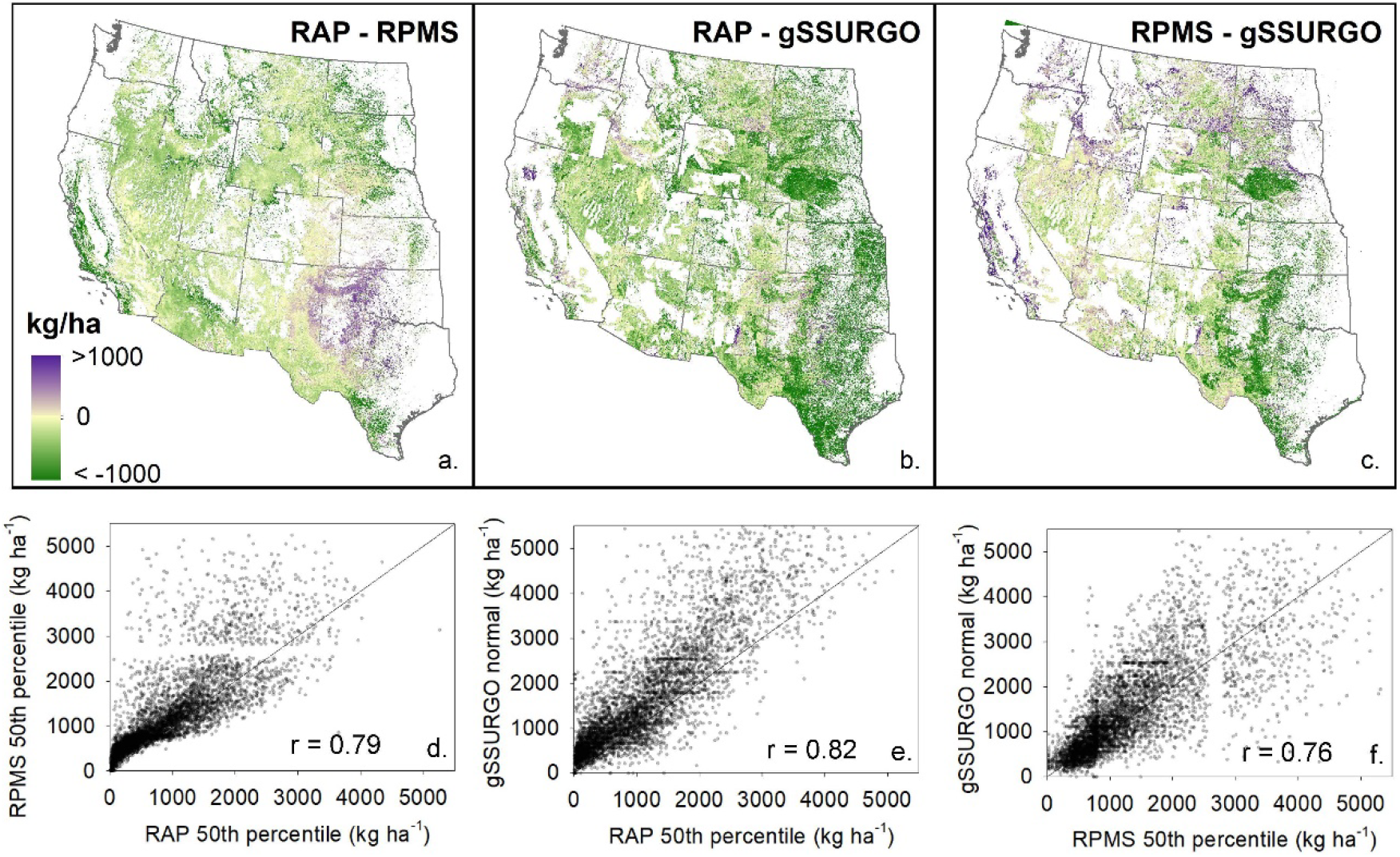
Differences (a-c) between three gridded production datasets across western U.S. rangelands using the 50th percentile of annual values (2000-2018) from RAP and RPMS data, and ‘normal’ values from gSSURGO. Scatterplots and Pearson correlation coefficients (d-f) of production values sampled from each data set for 5000 randomly selected rangeland locations.

Geographic distribution of the differences between the NRI herbaceous biomass estimates and the gridded HAGB, RPMS, and gSSURGO estimates (Supplemental Information) demonstrate general agreement across the western U.S. with greater differences apparent in the southern Great Plains.

## Discussion

Estimates of rangeland production–specifically herbaceous aboveground biomass or forage–are now available annually and at 16-day intervals from 1986 to 2019 at 30m resolution across the western U.S. and represent five advancements specifically relevant to management. These data are 1) provided in units recognized by and applicable to management (i.e., kg ha^−1^ or lbs acre^−1^); 2) produced at temporal fidelities applicable to monitoring the effects of climate, disturbance, management, and other factors, 3) calculated at a spatial resolution (30m) that allows for assessment of variability both within and across management units; 4) are easily accessible where monitoring, analysis, and interpretation can be achieved without the need for specialized technical knowledge or skills; and 5) account for annual pixel-scale changes in PFT composition (e.g. annual grasses or trees encroaching rangelands) and the relative contributions of annual forbs and grasses and perennial forbs and grasses to total herbaceous biomass. The availability of these data and other rangeland wide vegetation data (Reeves et al., 2020, Rigge et al., 2020, Pastick et al., 2020) have ushered in a new era where rangeland mapping from national to management scales is now a working reality (Jones et al., 2020).

These herbaceous biomass data provide land managers, practitioners, and decision-makers novel, temporally continuous, and spatially contiguous data for enhanced rangeland management. These data can be paired with local knowledge and information to better inform management strategies at the scale of a grazing allotment or pasture. At larger scales, these biomass data shed light on persistent ecosystem threats like invasive annual grasses (Jones et al., 2020) that are contributing to the continual rise of annual grass/forb biomass on western rangelands (Fusco et al., 2019; see Fig. 1). Differences in forage quality and phenology between annuals and perennials also have important implications for rangeland functions including accelerated wildfire return intervals (Pilliod et al., 2017). These represent only a few of the many potential scenarios where such data can provide greater insight and better inform management of grazing, wildlife, fuel, and fire, as well as assessing outcomes of management practices.

This technical note does not present a traditional validation of these new RAP HAGB data due to the lack of plot-level HAGB data at the scope and scale of the data product. While NRI plot-level data do include *some* destructive sampling (i.e., clipping, drying, and weighing), the methods also incorporate subjective estimations and correction factors. Also, the model used to estimate RAP HAGB is process based and not empirical – it does not incorporate biomass field plots at any step. We therefore examine ‘multiple lines of evidence’ and factorially compare the four available broad-scale data sets of rangeland production. This method demonstrates a best-practices approach when using these type of data in a decision-making framework; utilize all data sources, examine their similarities and discrepancies, and incorporate local knowledge to best inform a data driven decision.

## Implications

The temporally dynamic herbaceous aboveground biomass data represent a culmination of advancements in utilizing remote sensing data to monitor rangelands more effectively and efficiently. The new geospatial datasets of rangeland production provide land managers and decision-makers spatially contiguous and temporally relevant data to monitor rangeland forage, conduct meaningful comparisons of management outcomes using common data, and examine within season variability of forage to better assess management actions. The readily available data (RAP, https://rangelands.app/; RPMS, https://www.fuelcast.net/) remove analytical and technological barriers, allowing for immediate utilization. Never before has so much data been directly available and applicable to rangeland management. We anticipate and look forward to new applications, analyses, discoveries, and innovations with these data that improve our understanding and management of rangelands.

## Acknowledgments

This work was made possible by the United States Department of Agriculture, Natural Resources Conservation Service’s Working Lands for Wildlife, the USDA’s Conservation Effects Assessment Project, and the Bureau of Land Management. All data are freely available via the Rangeland Analysis Platform (https://rangelands.app).

