## Supplemental Information and Figure S1. for "Annual and 16-day rangeland production estimates for the western United States"

The temporal and spatial scale of the Rangeland Analysis Platform (RAP) herbaceous aboveground biomass (HAGB) estimates (as well as the use of a process-based light-use-efficiency model for calculating such estimates) inhibits the ability to conduct a formal validation. The Natural Resources Conservation Service (NRCS) National Resources Inventory (NRI) plot-level herbaceous biomass data (NRCS, USDA, 2015) provide the only field-level data at a similar geographic and temporal scale, but the methods incorporate subjective estimations and correction factors so must also be considered estimates. Nonetheless, viewing the geographic distribution of differences between plot-level estimates and gridded biomass products can be useful.

Figure S1 displays the geographic distribution of differences between the NRI plot-level estimates of herbaceous production and three gridded datasets. Differences at each plot location were calculated between the RAP HAGB estimates, the United States Forest Service Rangeland Production Monitoring Service (RPMS) annual data (provided annually from 1984-2018 at 250m resolution; Reeves et al., 2020), the gridded Soil Survey Geographic (gSSURGO) normal potential production data (Soil Survey Staff, 2017), and 16,591 Natural Resources Conservation Service (NRCS) National Resources Inventory (NRI) plot-level estimates of herbaceous biomass collected on rangelands from 2004 to 2018 (NRCS, USDA, 2015). The biomass estimates from RAP HAGB and RPMS were sampled from the same year as the plot measurement; gSSURGO data is temporally static. Pearson correlation coefficients between NRI estimates and gridded estimates were also calculated.

Greater differences between NRI estimates and gridded estimates occur in typically higher biomass areas such as the Great Plains and coastal California. It is important to note that although geographic patterns appear to be present, close examination reveals that differences in one location are not necessarily characteristic of the differences in other close-proximity locations, nor are differences homogenous at ecoregion scales. For example, the Flint Hills ecoregion (Figure S1 d,e,f) contains positive and negative differences within all three comparisons. Considering the gridded datasets are calculated using inherently different empirical and process-based models, and that plot-level NRI estimates contain subjective estimations and correction factors and are collected at a far different scale than the RAP HAGB gridded estimates (0.89 $m^2$ quadrats vs. 900 $m^2$ pixels), it is critical to not judge the efficacy of any one product based on these comparisons. Indeed, these differences reinforce a best-practices approach when using such data in a decision-making framework; utilize all data sources, examine their similarities and discrepancies, and incorporate local knowledge to best inform a data driven decision.

Soil Survey Staff, 2017 Natural Resources Conservation Service, United States Department of Agriculture. Web Soil Survey.

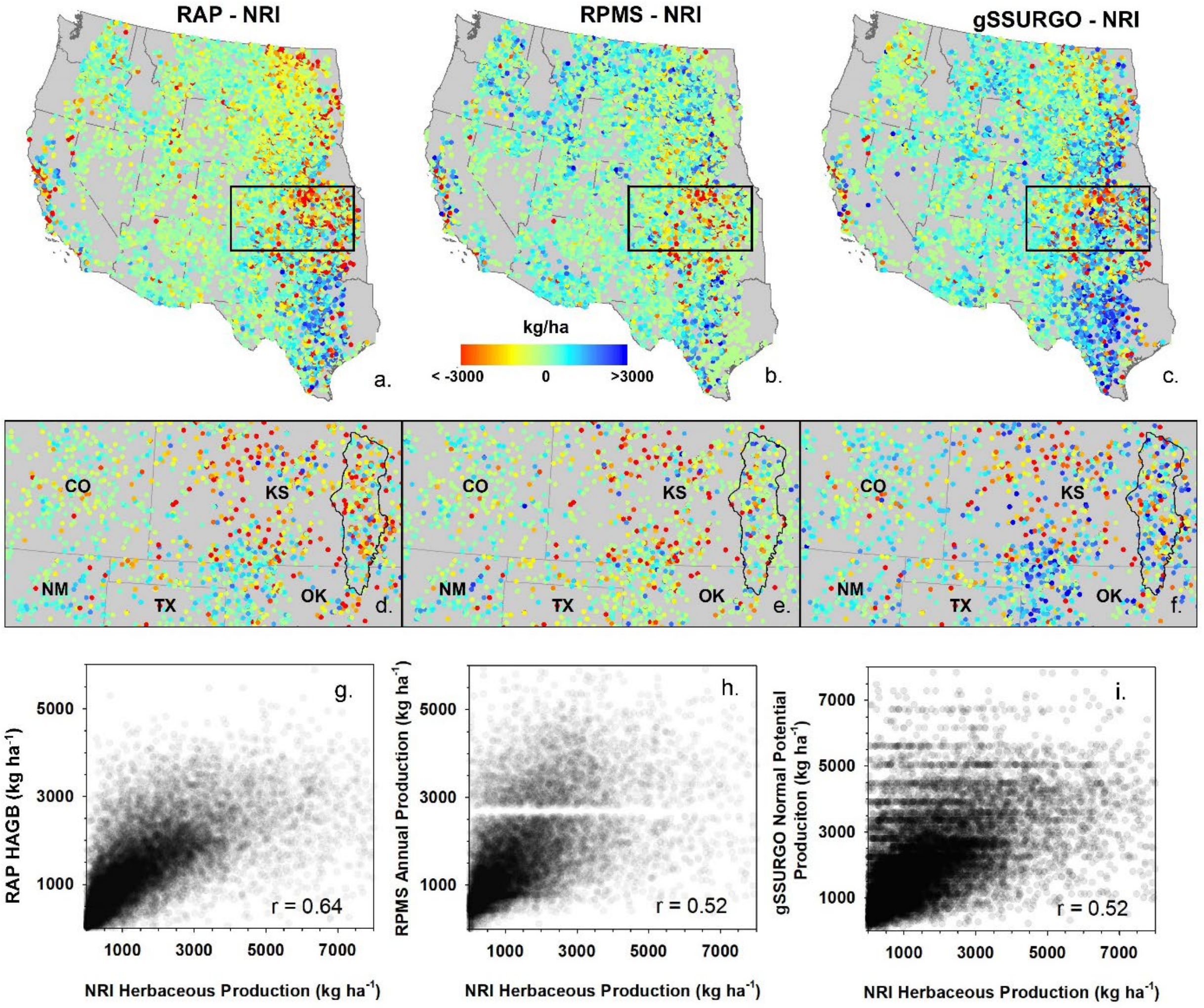


Figure S1. Difference between NRI plot-level estimates of herbaceous production and estimates from the Rangeland Analysis Platform herbaceous above ground biomass (HAGB) (a,d), Range Production Monitoring Service (RPMS) annual production (b,e), and gSSURGO normal potential production (c,f) gridded datasets. Black boxes in a, b, and c are extent of maps (d,e,f) which include state labels and the Flint Hills Level IV Ecoregion (black polygon). In all maps (a-f) values are NRI estimates subtracted from the gridded data. For the two temporally variable datasets (RAP HAGB, RPMS) the gridded data were sampled from the same year as the plot measurement. Scatterplots (g,h,i) display NRI plot-level estimates and corresponding value of gridded datasets. Pearson correlation coefficients (r-values) are presented within each scatterplot.
